# Human shape perception spontaneously discovers the biological origin of novel, but natural, stimuli

**DOI:** 10.1101/2024.12.21.629735

**Authors:** Kira I. Dehn, Guido Maiello, Frieder T. Hartmann, Yaniv Morgenstern, Sara Joy Hawkins, Thomas Offner, Joshua Walter, Thomas Hassenklöver, Ivan Manzini, Roland W. Fleming

## Abstract

Humans excel at categorizing objects by shape. This facility involves identifying shape features that objects have in common with other members of their class and relies—at least in part—on semantic/cognitive constructs. For example, plants sprout branches, fish grow fins, shoes are molded to our feet. Can humans parse shapes according to the processes that give shapes their key characteristics, even when such processes are hidden? To answer this, we investigated how humans perceive the shape of cells from the olfactory system of *Xenopus laevis* tadpoles. These objects are novel to most humans yet occur in nature and cluster into classes following their underlying biological function. We reconstructed 3D cell models through 3D-microscopy and photogrammetry, then conducted psychophysical experiments. Human participants performed two tasks: they arranged 3D-printed cell models by similarity and rated them along eight visual dimensions. Participants were highly consistent in their arrangements and ratings and spontaneously grouped stimuli to reflect the cell classes, unwittingly revealing the underlying processes shaping these forms. Our findings thus demonstrate that human perceptual organization mechanisms spontaneously parse the biological systematicities of never-before-seen, natural shapes. Integrating such human perceptual strategies into automated systems may enhance morphology-based analysis in biology and medicine.

## INTRODUCTION

What makes a shoe a shoe and not a fish? Whenever we look at a novel object, we effortlessly recognize what group or class it belongs to based on our experience with previous objects. This facility involves identifying key features of the object’s 3D shape that it has in common with other members of its class. These shared shape features are likely due to the shared processes that generated the objects in the first place. For example, plants sprout branches and leaves, fish grow a tails and fins to propel them through water, shoes are molded into a shape that matches the structure of human feet. Given that humans are able to parse shape features according to their physical causes^1–7^, this suggests that we may be able to rely on visual 3D shape perception to intuit the shared origins of objects. Yet these abilities rely—at least in part—on cognitive and semantic processes and constructs^8^. What happens instead when we are presented with completely novel objects of unknown origin? Remarkably, a wealth of previous research has demonstrated that humans can classify novel objects from few samples^9–13^. For example, Morgenstern et al.^11,12^ have shown that humans perform such “one-shot” categorization of novel object classes by relying both on simple heuristics as well as deeper analyses of shape characteristics, which may involve inferring the causal origin of an object’s form. However, most of this previous research has employed artificially generated stimuli, where the generation processes were constructed ad-hoc by the experimenters. This is problematic, as it pre-assumes which shape features make up a class. Yet natural objects can have extremely complex causal origins, leading to an almost unlimited range of possible diagnostic features for organizing them into classes. In this context, biological entities make particularly fascinating objects of perception, because they have richly structured shapes that result from exquisitely complex morphogenetic processes. It thus remains unknown whether the human visual system is able to recover the underlying biological origin of objects it has never seen before. This question is challenging to answer however, since humans have extensive experience with both natural and artificial generative processes in our everyday environment. We thus turned to an environment alien to most: the microscopic realm of cells (**Figure 1**).

**Figure 1.**
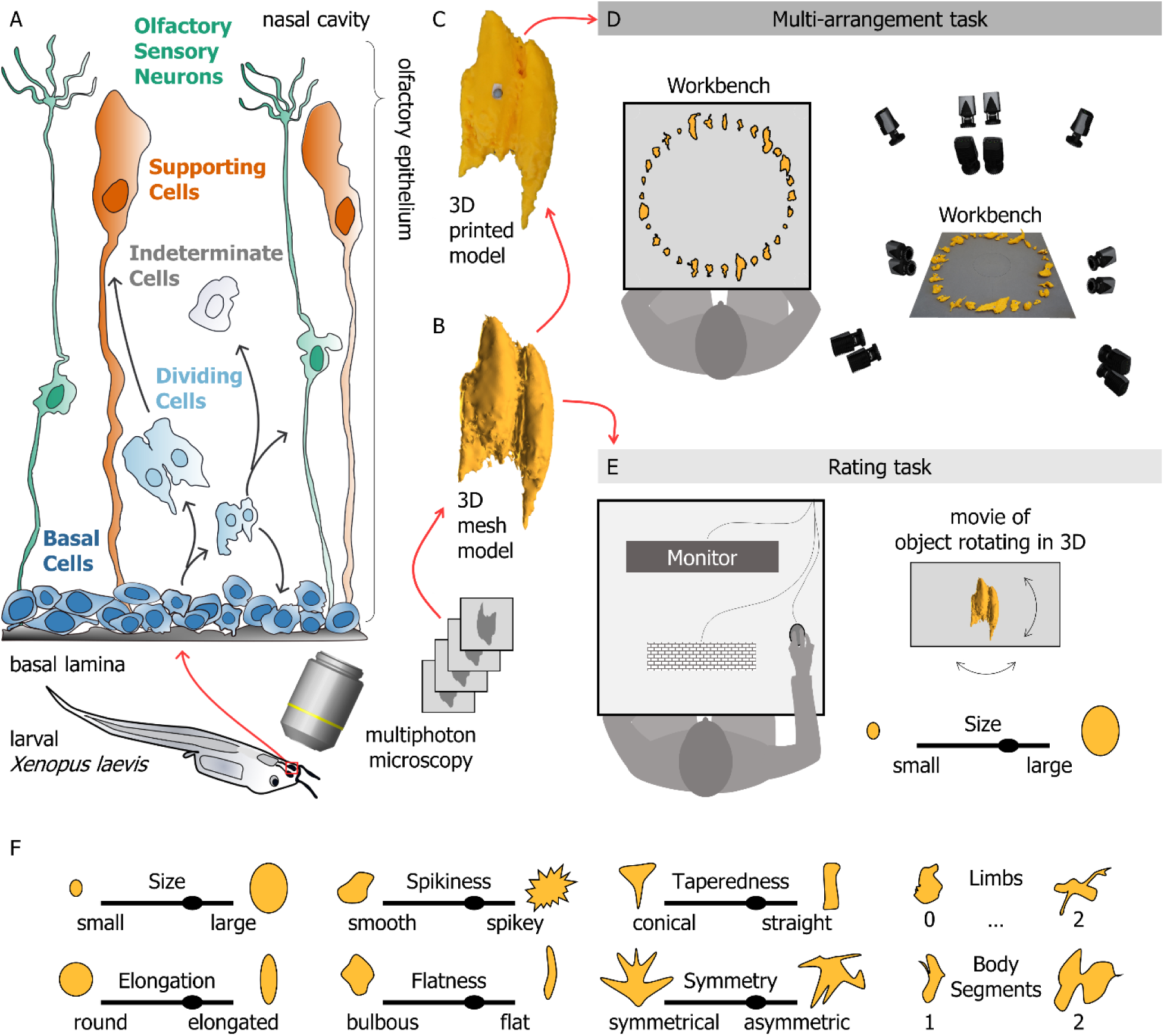
Using novel but natural stimuli to probe human 3D shape perception. (**A**) Experimental stimuli were 3D reconstructions of cells from the olfactory epithelium of Xenopus laevis tadpoles. The cell types reconstructed included basal cells, dividing cells, olfactory sensory neurons, supporting cells, and a set of indeterminate cells that could not be decisively assigned to any of the previous types. We reconstructed 30 individual stimuli, 6 stimuli per class. (**B**) For each of these cell types, 3D mesh model reconstructions were obtained through 3D-microscopy. (**C**) These 3D mesh models were then 3D printed, and a spherical retroreflective marker was glued onto each object. (**D**) During the multi-arrangement task, participants were seated at a workbench imaged from multiple angles using passive marker tracking cameras. The experimenter placed the 3D printed stimuli in random order in a circle on the workbench. Participants were asked to re-arrange the stimuli according to their shape similarity. The final object positioning was reconstructed and recorded using a position tracking system. (**E**) In the rating task, participants viewed stimulus videos on a computer monitor. Videos were renderings of each object rotating in depth. Participants rated the stimuli along each of eight hand-selected feature dimensions, specified in (**F**).

More specifically, we investigated how humans perceive and parse the 3D shape of cells from the olfactory epithelium of tadpoles of the African clawed frog *Xenopus laevis*^14^. These cells belong to different classes depending on their biological purpose (**Figure 1A**). The olfactory epithelium contains three main cell types: *basal cells*, *supporting cells* and *olfactory sensory neurons*^15^. Basal cells—found close to the basal lamina—are central to regenerative processes^16–18^ in the vertebrate olfactory system and may thus undergo mitosis (i.e. cell division). These *dividing cells* may then develop into either olfactory sensory neurons or supporting cells. Immature cells that have just undergone mitosis and have yet to develop clearly distinguishing features are classed as *Indeterminate*. To obtain 3D reconstructions of cells belonging to each of these classes, we acquired two-photon microscopy images of individual cells^19–21^ and input these into 3D photogrammetry software u-shape3D^22^, that created 3D mesh models of each cell (**Figure 1B**). We then 3D printed these 3D cell models at a scaling factor of 1:1000 (**Figure 1C**). Finally, we used these 3D printed objects and 3D renderings as stimuli in human behavioral experiments (**Figure 1D**,**E**).

In order to classify the cells according to their underlying classes, biologists often visually assess microscopy images of the cells^23^. These visual classifications primarily rely on identifying the stereotypical morphology of each cell class^14,24^. For example, basal cells tend to be small, round, and bulbous. Supporting cells and olfactory sensory neurons are instead elongated as they extend from the basal lamina toward the nasal cavity. Further, olfactory sensory neurons have stereotypical axonal and dendritic projections, whereas supporting cells exhibit a conical shape related to their structural purpose^14,15^. Dividing cells are often easily identifiable as they appear to be made of two body segments. Such manual, visually-based classifications can be an important in step in neurobiological investigations, yet they are time consuming and prone to human error^25^. It would thus greatly benefit neurobiological science if the process of morphology-based cell classification could be improved and/or automated^26^.

The purpose of this study was thus twofold, spanning two very distinct disciplines. Our first aim was to test whether human perceptual organization mechanisms parse 3D shapes according to the processes that give shapes their key characteristics, even when these processes are completely hidden. Additionally, we sought to leverage this human facility to develop and improve morphology-based cell classification systems. To achieve these goals, we designed two behavioral tasks. We designed a multi-arrangement task^27^ (**Figure 1D**) in which we asked participants to arrange 3D printed objects by how similar they appeared in shape. This task was meant to reveal how naïve observers would group novel stimuli, and whether such groupings might reflect the biological origin of the stimuli. We then designed a rating task (**Figure 1E**) which probed eight hand-selected visual feature dimensions (**Figure 1F**). This tested whether we could define ad-hoc shape features that were related to human similarity judgments and that could also be used to differentiate the cell classes.

## RESULTS

### Experiment 1 – Multi-arrangement task

In Experiment 1, we used a multi-arrangement task (**Figure 1D**) to investigate whether naïve observers would spontaneously group novel stimuli, and whether such groupings might reflect the biological origin of the stimuli. We asked participants to arrange 3D printed objects by similarity. All participants were tested with the same set of 30 stimuli (6 per class). Participants performed this multi-arrangement task once at the beginning of the experiment (1^st^ session) and once at the end (2^nd^ session). Between these two sessions, participants performed a rating task (**Figure 1E**, discussed below).

#### Participants spontaneously grouped novel stimuli, and groupings were related to the biological cell classes

To visualize participant arrangements, we aligned multi-arrangement data using Procrustes analysis. Specifically, we rigidly translated and rotated the stimulus arrangements so they would match as closely as possible across sessions and participants, regardless of their orientation on the workbench. Upon first visual inspection, how participants arranged the stimuli seemed consistent across sessions and broadly corresponded to the underlying ground truth cell classes (**Figure 2A**). To measure how similarly participants grouped the stimuli, we computed representational dissimilarity matrices (RDMs)^27^ for each session and for every participant. RDMs were constructed by computing pairwise Euclidian distances between every possible stimulus combination, normalized to the maximum distance between stimuli. The patterns of correlation between RDMs across sessions and participants represent the agreement between these different arrangements. The RDMs from one example participant (**Figure 2B**) illustrate the high internal consistency across sessions (r=.94, p<.001). The similarity of the example participant’s RDMs to the group average RDM (**Figure 2C**, left) also illustrates how different participants produced similar arrangements. These robust within- and between-participant agreements held true across our sample (**Figure 2D**, within-participant r=.87, p<.001, between-participant r=.66, p<.001).

**Figure 2.**
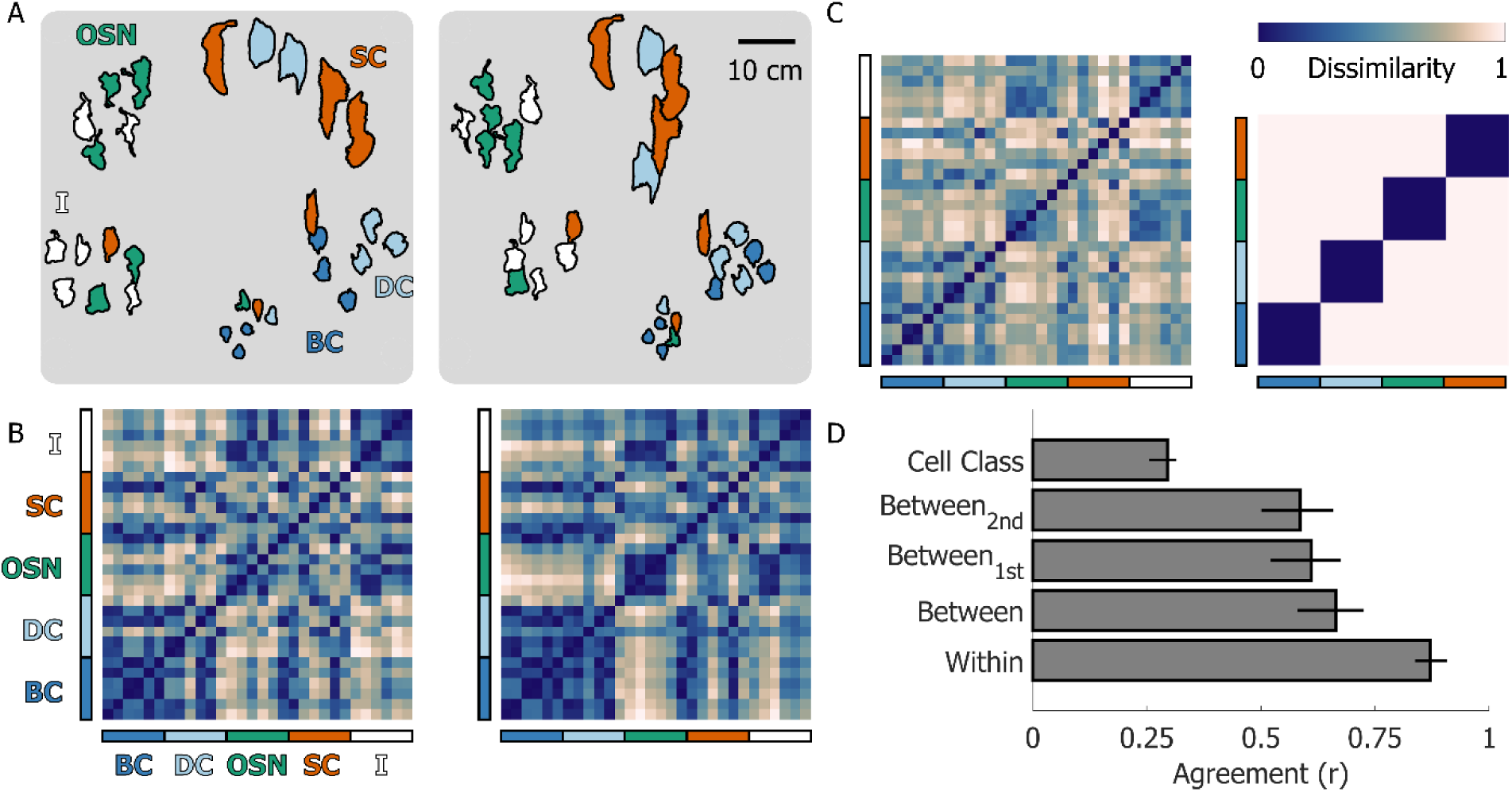
Multi-arrangement results. (**A**) Multi-arrangement data for one example participant in the 1^st^ (left) and 2^nd^ sessions (right) of the multi-arrangement task from Experiment 1. Data from the two sessions were aligned using Procrustes analysis. Different colors represent different cell classes: (BC) Basal cells; (DC) Dividing cells; (OSN) Olfactory sensory neurons; (SC) Supporting cells, (I) Indeterminate cells (**B**) Representational dissimilarity matrices (RDMs) for the 1^st^ (left) and 2^nd^ sessions (right) from the same example participant. Dissimilarity was defined as the normalized Euclidian distance between stimulus pairs. Colored bars along the x- and y-axis of the RDMs color-code the cell classes. (**C**) Mean RDM averaged across participants and sessions (left) compared to the ground truth classification RMD derived from the cell classes. Note that we excluded Indeterminate cells, as these may belong to different biological cell classes. (**D**) Within and between-participant agreement in the multi-arrangement task and agreement with the ground truth cell classes. Bars represent the mean across participants; error bars represent the 95% bootstrapped confidence intervals of the mean.

Individual participants thus relied on specific strategies when arranging the stimuli, and these strategies were similar across participants. This suggests that there may be a common set of features that participants employed to judge similarities between 3D shapes. These features might in turn be related to the biological origin of the stimuli. If this were true, participants should have grouped the stimuli lawfully according to their underlying cell classes. To test this, we computed the dissimilarity matrix for perfect ground truth classification and compared it to the average RDMs of all participants from Experiment 1. As shown in **Figure 2C**, the group average participant RDM (left) exhibits some—albeit weak—structure along the diagonal which aligns with the cell class RDM (right), giving rise to the weak but significant correlation between participant arrangements and biological cell class (**Figure 2D**, r=.29, p<.001). This correlation was clearly not as strong as the between-subject correlation, but this is possibly because the pure cell class RDM was not designed to capture the patterns of similarity across cell classes.

Between the 1^st^ and 2^nd^ multi-arrangement sessions, participants preformed a 30-minute rating task in which they rated the same stimuli along a set of visual feature dimensions. We wondered whether the rating task might have induced participants to change strategies between sessions and arrange the stimuli according to the same set of rated dimensions. If this were the case, we might expect between-participant agreement to be stronger in the 2^nd^ session compared to the 1^st^ session. Therefore, we calculated between-participant agreement for the 1^st^ (r=.61, p<.001) and 2^nd^ sessions (r=.58, p<.001) separately (**Figure 2D**) and observed that between-participant agreements did not differ significantly across sessions (*t*(15)=.975, p=.345). We thus found no evidence that the rating task influenced participant arrangement strategies.

### Experiments 1 and 2 – Rating task

We designed a computer-based rating task which probed eight hand-selected visual feature dimensions. Participants viewed movie renderings of individual 3D shape stimuli and rated each stimulus along each dimension. Stimulus objects were the same as those employed in the multi-arrangement task. We selected the feature dimensions to span a perceptual space that would plausibly cover the 3D shape characteristics of the cell stimuli. In Experiment 1, participants performed this rating task between the 1^st^ and 2^nd^ multi-arrangement sessions. We tested whether participants could reliably rate the stimuli along these feature dimensions, and whether performance at the rating task was as reliable as participant arrangements. We also assessed whether feature ratings were related to how participants arranged the stimuli in the multi-arrangement task. In Experiment 2, we verified whether a different set of participants could perform feature ratings through vision alone, i.e., without having ever touched or manipulated the 3D objects. Finally, we used Principal Component Analysis (PCA) to examine the structure of the feature space and tested whether the selected feature dimensions were related to the ground truth biological cell class.

#### Participants could reliably rate novel stimuli along hand-selected feature dimensions

**Figure 1F** lists the eight hand-selected feature dimensions we investigated. Participant ratings qualitatively aligned with the feature dimensions. For example, the stimuli increase in size if we organize them according to average participant “Size” ratings (**Figure 3A**). If instead we organize the stimuli according to the average participant “Spikiness” ratings, we can observe how the surface texture visually changes from smooth to spikey (**Figure 3B**). Further, participants agreed with one another on each of the feature dimensions (**Figure 3C**), meaning that different participants gave similar ratings to each of the stimulus objects. Indeed, the between-participant level of agreement for 7/8 feature dimensions was as good as or better than the between-participant agreement in the multi-arrangement task.

**Figure 3.**
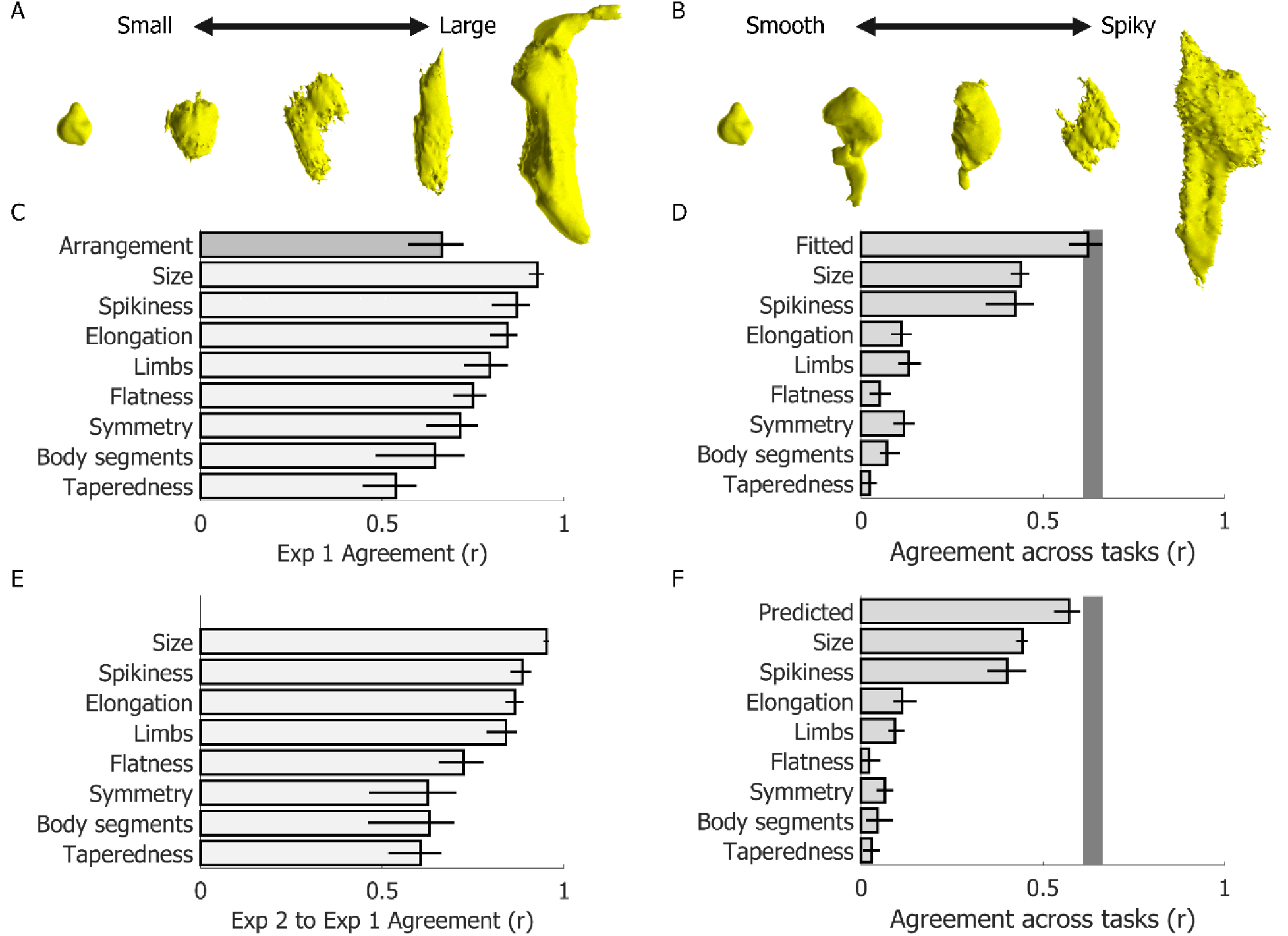
Between-participant agreement within and across experiments and tasks. (**A,B**) Stimuli ranked 1^st^, 7^th^, 15^th^, 22^nd^, and 30^th^ from the average participant ratings of the “size” and “spikiness” dimensions. (**C**) Between-participant agreement for each of the rating dimensions in Experiment 1. As reference, the darker grey bar displays the between-participant agreement for the same participants in the multi-arrangement task. (**D**) Agreement between rating and multi-arrangement data in Experiment 1. (**E**) Agreement between participant ratings across Experiments 1 and 2. (**F**) Agreement between rating and multi-arrangement data across Experiments 1 and 2. Bars represent the mean across participants; error bars are the 95% confidence intervals of the mean. Grey shaded region in D,F represents the noise ceiling (i.e., the upper and lower estimates of the between participant agreement in the multi-arrangement task, corresponding to the upper bound of possible agreement across tasks).

#### Perceptual ratings almost perfectly predicted how participants arranged stimuli in the multi-arrangement task

To assess whether feature ratings were related to how participants arranged the stimuli in the multi-arrangement task, we computed RDMs for each rating dimension and for every participant. We then correlated these RDMs with the average RDM from the multi-arrangement task. We further fit a linear model regressing individual participant rating RDMs onto the average multi-arrangement RDM. **Figure 3D** shows that “size” and “spikiness” are the dimensions that best explain the participant arrangements. It is perhaps interesting to note that these were also the rating dimensions that participants agreed on most strongly. Further, a simple linear combination of the feature ratings could reliably explain participant arrangements. Indeed, the best-fitting model correlated with participant arrangements as strongly as the between participant agreement in the multi-arrangement task (which here we take as a measure of the noise ceiling). We cross-validated the model using a different set of participants in Experiment 2 (see below).

#### Participants could rate the stimuli using vision alone

We wondered whether haptically manipulating the printed 3D models could have influenced participant behaviour in the rating task. To test this, we thus ran a second Experiment. We asked a separate set of participants to perform only the feature rating task, without ever physically interacting with the physical 3D stimuli. We then correlated the rating data of individual participants from Experiment 2 to the average ratings across participants from Experiment 1. The strong and significant correlations between participants and across experiments (**Figure 3E**) suggest that the observers’ behaviour on the rating task was not influenced by the multi-arrangement task. Further, we took the coefficients of the model fitted to the Experiment 1 data, which were based on ratings from Experiment 1, and applied them to participant ratings from Experiment 2. This provided cross-validated predictions of participant arrangements from Experiment 1. These predictions could explain participant arrangements from Experiment 1 nearly as accurately as the participant ratings from Experiment 1 itself (**Figure 3F**). This suggests that visual shape perception was sufficient to explain how participants arranged the stimuli in Experiment 1.

#### Rating data spanned a multidimensional space that linearly separated objects into their underlying cell classes

We next tested to what extent the rating dimensions we chose were independent and examined the structure of the perceptual space they spanned. **Figure 4A** shows overall moderate-to-low correlations between the feature dimensions, whereas **Figure 4B** shows that nearly all eight PCA dimensions are necessary to account for the variance in the rating data. These results suggest that the feature dimensions we selected were at least partly independent. We next fit a multiple linear regression model to attempt to predict cell class from the PCA transformed rating data. The fitted model significantly correlated with cell class, also slightly better than the multi-arrangement data (**Figure 4C**). Indeed, cell class appeared to be visually separable when projected onto the first two PCA dimensions (**Figure 4D**). Further, cell class in PCA space appeared to be more separable but less clustered than in the multi-arrangement data (**Figure 4E**), which explains why the correlation between feature rating data and cell class was still relatively low. However, these patterns suggested that a linear classifier should be able to tease apart cell class from both rating and multi-arrangement data.

**Figure 4.**
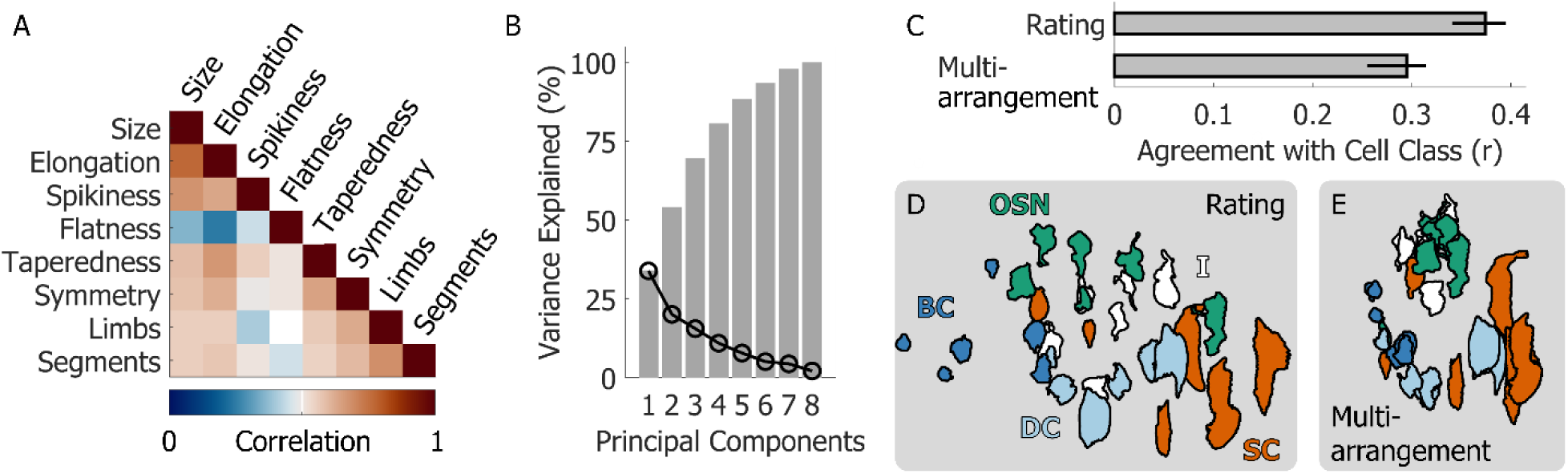
The structure of the rating data. (**A**) Multicollinearity matrix showing the correlations between the eight feature dimensions probed by the rating task. (**B**) Scree plot displaying the variance explained (black curve) and cumulative variance explained (grey bars) the Principal Components of the rating data. (**C**) Agreement of the multi-arrangement and rating data with the ground truth cell classes. (**D**) The average rating data projected in the first two Principal Components separate the known cell classes. (**E**) The average multi-arrangement data are more clustered but do not separate cell class as clearly.

### Experiments 1 and 3 – Recovering cell class from multi-arrangement vs rating data in naïve observers and experts

One of the goals of this project was to test whether we could leverage human visual 3D shape perception to classify cell stimuli into their underlying biological classes. We were also interested in whether participants need to be familiar with the underlying cell classes to provide data that could tease these classes apart. For this reason, we ran a final Experiment 3 in which we asked a group of five expert biologists to perform the same tasks performed by naïve participants in Experiment 1. In previous analyses cell class appeared to be—at least to some extent—linearly separable from the multi-arrangement and rating data. We therefore employed support vector machine classifiers to attempt to recover cell class from multi-arrangement and rating data in both naïve participants and experts.

#### Support vector machine classifiers recovered cell class across tasks and participant groups

We employed a cross-validation procedure to gauge whether support vector machine classifiers could predict cell class from multi-arrangement and rating data provided by naïve participants (Experiment 1) and expert biologists (Experiment 3). For each data type and participant group, we trained classifiers on 20 cell stimuli (5 stimuli per each cell class, excluding the indeterminate cells) and tested whether the classifiers correctly predicted the cell class of the four left-out cells. We repeated this procedure for all possible combinations of four left-out cells. Each iteration, we also queried the classifiers to predict the class of the indeterminate cells.

The average human data, projected in 2D space, appeared qualitatively similar across tasks and participant groups (left subpanels of **Figures 5A-D**). Cells belonging to the same class were near one another, and the spatial relations between cell classes were preserved. For this reason, cross-validated classification accuracy was well above chance across tasks and participant groups (**Figure 5E**). The cross-validated confusion matrices (right subpanels of **Figures 5A-D**) show which cell classes were most easily confused with one another. Basal cells (dark blue) and olfactory sensory neurons (green) were rarely misclassified. Dividing cells (light blue) and supporting cells (orange) were instead more easily confused with each other. This makes sense from a biological viewpoint, as dividing cells may be turning into supporting cells. Further, across all datasets the indeterminate cells (white) were predominantly classified as olfactory sensory neurons. This suggests that in our stimulus set the category of the indeterminate cells consisted predominantly of immature olfactory sensory neurons, which already displayed shape features similar to those of the mature cells, such as the characteristic axonal and dendritic protrusions. This possibility is corroborated by the fact that expert biologists grouped 5/6 of the indeterminate cells with the olfactory sensory neurons in the multi-arrangement task (**Figure 5C**).

**Figure 5.**
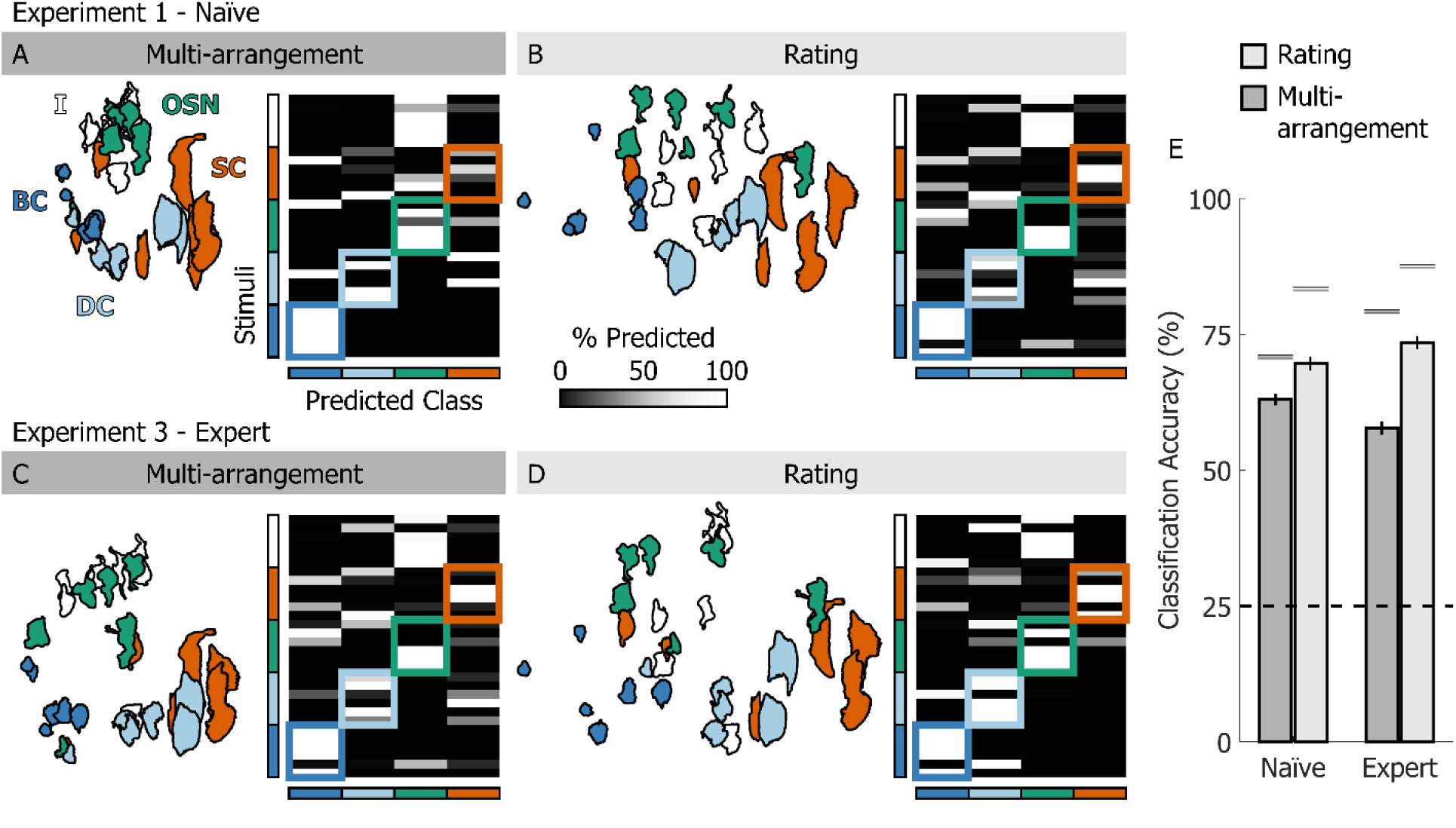
Classification analyses. (**A**) Left: multi-arrangement data from naïve participants from Experiment 1, averaged across participants. Right: cross-validated confusion matrix for a Support Vector Machine trained to classify cell class from the multi-arrangement data. (**B**) Left: rating data from naïve participants from Experiment 1, projected in PCA space and averaged across participants. Right: cross-validated confusion matrix for a Support Vector Machine trained to classify cell class from the PCA-projected rating data. (**C,D**) As A,B, except for expert biologists from Experiment 3. (**E**) Cross-validated classification accuracy for all SVM classifiers trained on different data and participants. Bars are means, error bars are 95% confidence intervals. Dashed line represents chance performance; grey lines show best-achievable classification accuracy.

For both experts and naïve participants, cross-validated classification accuracy for the classifiers trained on the rating data was higher compared to the classifiers trained on the multi-arrangement data (**Figure 5E**). This was in part due to the higher dimensionality of the rating data. However, this pattern held true when we performed the same analyses using only the first two PCA dimensions of the rating data. Furthermore, it is striking that the classification accuracy obtained from the naïve participants’ data was nearly as high as that obtained from the experts’ data. These results suggest that a feature rating task may be preferable to a spatial arrangement task for classifying cell stimuli into their underlying biological classes. Further, no knowledge of the origin of the stimuli is required to rate visual shape features that can distinguish the different cell classes.

## DISCUSSION

Humans are able to visually identify and categorize objects based on, among other things, the objects’ perceived 3D shape^1–13^. Here, we investigated this ability by using a unique set of stimuli that have a common biological origin but are unknown to most: 3D models of cells from the olfactory system of *Xenopus laevis* tadpoles. We asked human participants to manually sort 3D printed versions of these stimuli and rate animated renderings along a set of predetermined feature dimensions. Human arrangements and ratings were highly reliable within and across participants and tasks. Participants’ arrangements clustered systematically, and indeed biological cell class was linearly decodable from both arrangement and rating data.

### Can humans leverage generative models to infer the causal origin of never-before seen shapes?

Previous studies have demonstrated that humans are able to make precise categorization decisions based on a small number of examples^9–13^. Such “one-shot” categorization of novel object classes may involve inferring the causal origin of the objects’ form^11,12^. However, previous research has demonstrated that these abilities may rely in part on cognitive and semantic processes and constructs^8^. For example, Spröte et al.^4^ used both familiar and unfamiliar 2D shapes that appeared to be “complete” or “bitten” and asked participants to place dots to identify their perceived symmetry axis. Participant responses for “complete” and “bitten” objects were similar, suggesting participants could identify and discount the process that generated the object deformations (i.e. the “biting”). To study object categorization independently of such higher-level cognitive constructs (such as the notion of an object part being “bitten-off”), some researchers have employed discriminative tasks with purely artificially generated stimuli and artificially designed generative processes^28^. Discriminative tasks such as object categorization however have a crucial problem when it comes to pre-selected stimuli. The experimenter determines the categories or, in the case of artificial stimuli, the relevant features and/or generative processes that determine “ground truth” category labels. Then, participants typically compare artificial objects to one another, thus constraining the investigations to the experimenters’ expectations—reflected in the selected features and/or artificial generative processes.

Tiedemann et al.^13^ in part overcame these limitations using a drawing task in which participants viewed 2D “Exemplar” shapes and then created their own “Variations” of these shapes belonging to the same class. The participant-drawn “Variations” were not mere copies of the “Exemplars” yet were strikingly similar across participants and shared distinctive features. Participants thus behaved as if they were employing common generative models to parse the “Exemplars” into categories. However, even in this study the “Exemplars” were artificially generated by the experimenters, and the results were potentially constrained by the participants’ drawing skills. These findings could thus still reflect higher-level cognitive processes shared between participants and experimenters.

Our experimental design addressed this potential issue by asking participants to arrange— according to the perceived 3D shape similarity—novel real-world objects which fell into unbiased ground truth categories. In line with Tiedemann et al.^13^, here we found that participants produced robust and consistent arrangements internally as well as between each other. Additionally, participant judgments broadly agreed with the underlying object classes, even though we did not explicitly ask participants to uncover such classes. Given that these classes arose from different natural generative processes, our findings corroborate the notion that humans can recover such generative processes and use generative models to analyze novel shapes.

### A high dimensional shape feature space

Our results thus demonstrate that human can visually tease apart the underlying biological origin of never-before-seen stimuli. What substrate does this ability rely on? Previous literature has proposed that processes of perceptual organization, such as the decomposition of shapes into different parts according to their geometric properties^29–34^ are a key aspect of segmenting and representing shapes. Specifically, humans might represent objects as points in a high dimensional feature space in which objects that are close together belong to the same class, and objects that are far apart belong to different classes^35–38^. To operationalize this hypothesis, Morgenstern et al.^11^ developed ShapeComp, an image computable model— based on over 100 2D shape descriptors—capable of accurately predicting human shape similarity judgments. These authors then employed ShapeComp to study how humans categorize novel object shapes. When participants were asked to categorize novel objects into different classes, and were given multiple exemplars of each class, participants relied on a fixed set of ShapeComp features to perform the task. When participants were given just one exemplar per class instead (i.e. one-shot categorization), they relied on a more sophisticated strategy that involved flexibly re-weighting features on a stimulus-by-stimulus basis. This suggested that the feature space is more important than the specific features it is composed of. Our results are consistent with, and substantially extend, these previous findings obtained on 2D shape silhouettes derived from animal shapes. Notably, our arrangement task was neither a one-shot categorization task, nor could participants rely on multiple class exemplars as the stimuli were novel and we did not explicitly tell participants that the stimuli belonged to different classes. Nevertheless, we were able to define a set of ad-hoc visual features that were predictive of participant shape similarity arrangements. This set of hand-selected features was also sufficient to tease apart the stimulus classes. Therefore, our results demonstrate that our participants’ ability to organize novel stimuli according to their hidden underlying causal origin relies on a highly flexible feature space that can be reweighted to reflect the statistical properties of the stimulus set.

### Bringing together feature-based and generative models of shape perception

Spröte et al.^4^ suggest that the underlying generative processes are a key component to structure and organize perceived shape and a possible strategy to make assumptions of how other members of the same class might look. Fleming and Schmidt^28^ instead propose that it is the objects’ statistical features that are responsible for their correct identification, and not the generative process itself. Taken together with this previous research, our findings suggest that both could be true. Specifically, in our stimulus set the members of each cell class share a common growth process that likely accounts for the specific shape features shared by members of a same cell class. Mental generative models could thus operate directly in a shape feature space, by first uncovering the statistical feature regularities within a stimulus set, and then reweighting the feature space to reflect these regularities. Then, tasks requiring quick judgments of novel shapes (e.g. one-shot object categorization) could be performed by assessing the plausibility of items and trajectories within this reweighted shape feature space.

### Biological cell classification

Our findings relating to the visual analysis of cell shape may also be of interest to the research field of biological cell classification. To give just a few examples, cell shape guides the formation of healthy cardiac chambers^39^, dictates whether individual cells grow or die^40^, and can help predict disease progression^41^. The analysis of cell morphology is thus of great interest across biology and medicine, and although researchers have been developing automated methods to conduct cell shape analysis, most previous attempts have focused on 2D images^42,43^. The u-shape3D^22^ pipeline employed in this work to reconstruct 3D cell shapes from two-photon microscopy images of individual cells is one of the first tools for cellular 3D morphometry, but this field is still in its infancy. Our work suggests that leveraging the computational principles of human visual 3D shape processing may be a way forward. Specifically, simple machine learning approaches could be used to recover cell class from human shape feature ratings, or from stimulus-computable quantities that model these.

Examining the spatial organization of participant arrangements and ratings in **Figure 5**, is even more intriguing. For example, we can note that basal cells are adjacent to dividing cells, and dividing cells are in turn adjacent and overlapping with supporting cells. These patterns appear to reflect the underlying biology (**Figure 1a**), as dividing cells are in fact basal cells undergoing mitosis to replenish other cell types in the olfactory epithelium^14^. Olfactory sensory neurons are—for the most part—clustered away from other cell types. This is likely due to their characteristic axonal and dendritic protrusions, which make them saliently different from the other cell classes^15^. These protrusions develop only after cell division, which is likely why the dividing cells do not overlap with the olfactory sensory neurons. Further, the indeterminate cells seem to be located between supporting cells and olfactory sensory neurons. This reflects the fact that the indeterminate cells are likely immature cells developing into either supporting cells or olfactory sensory neurons. These observations reveal an even deeper insight: we may be able to leverage the principles of human 3D shape processing to map out the life cycle and developmental trajectories of cells within their environment.

### Why aren’t the data perfectly clustered?

In our study, participants had no prior knowledge of the biological processes that shaped the objects and were asked to arrange the stimuli purely based on their visual similarity. While the natural cell classes were not perfectly mirrored in the similarity arrangements, as we note above more nuanced patterns emerged in the data (**Figure 5**). This only partial alignment with biological classification is thus not surprising given that many of these cell classes share a common morphogenetic history. For example, dividing cells can develop into supporting cells or olfactory sensory neurons (**Figure 1A**). Such shared developmental pathways may have made it difficult for participants to fully differentiate between distinct cell classes based solely on shape. Moreover, when selecting the stimuli, we focused on cell class but not on the cells’ specific degree of maturity. Immature cells, or even cells in the process of dividing, may introduce ambiguity into the perceived shape features, making it more challenging to identify their function. This overlap in statistical features across developing or dividing cells likely blurred the boundaries between classes in the participant arrangements. Additionally, while experts performed slightly better, their data were not significantly more discriminative of cell class than the data from naïve participants. This may be due to the fact that experts, although trained to classify cells, were still arranging stimuli based on visual similarity, rather than explicitly grouping them into known biological categories.

### Future Directions

Our study opens up many avenues for future research. For example, further investigation is needed to determine the role of tactile manipulation and exploration^44,45^ in constructing internal generative models of 3D shape. Additionally, participants in our study might have been able to group stimuli more reliably according to their cell class if they had been explicitly asked to uncover these underlying groups. In particular, expert biologists usually use more than just shape features when classifying cells, by taking into account a cell’s location and orientation in the olfactory system to determine its type and function^14^. Incorporating these contextual cues in future experiments might improve classification accuracy for both experts and naïve participants. Further, biologists often observe cells through various imaging systems and in different environments, which likely influences their classification process in ways that were not captured by our task. Nonetheless, this study provides a proof of concept that human 3D shape perception can be leveraged to classify and map out different cell classes in biological systems.

Moving forward, several strategies could be employed to enhance this approach. First, crowdsourcing^46^ could be used to gather a broader set of perceptual data, possibly revealing additional common strategies for grouping unfamiliar biological shapes. Second, biologically inspired computer vision systems^47–52^ could be developed or adapted to replicate human strategies for parsing 3D shapes and distinguishing between subtle biological features. These systems could eventually surpass human capabilities in tasks like cell classification by incorporating contextual data or learning to recognize features that are not immediately obvious to the human eye.

In conclusion, by focusing on objects that exist in nature but with which humans have limited experience—i.e., cellular structures—we were able to demonstrate that human visual perception can successfully categorize novel 3D shapes based on their underlying biological origins. Our approach reveals the flexibility of the human perceptual system in organizing unfamiliar objects according to shape features, even in the absence of contextual information. These findings provide a new understanding of how humans visually process 3D shape and suggests practical applications in fields like biology and medicine, where leveraging human perceptual strategies could aid in automating tasks such as cell classification and morphological analysis.

## METHODS

### Participants

Participants were students and staff from Justus Liebig University Giessen. All participants had normal or corrected to normal vision and participated in the experiments for course credit or financial compensation (at a rate of 8 Euro/h). We recruited 16 naïve participants for Experiment 1 (12 female, mean [range] age 23 [18-27] years), 15 naïve participants for Experiment 2 (9 female, 26 [20-35] years), and 5 “expert” participants for Experiment 3 (4 female, 28 [25-32] years). Expert participants were current or past members of the Animal Physiology and Molecular Biomedicine research group at the Institute of Animal Physiology of Justus Liebig University Giessen. All participants provided written informed consent. All procedures were approved by the local ethics committee of Justus Liebig University Giessen and adhered to the tenants of the sixth revision of the declaration of Helsinki (2008).

### Stimuli

Experiments included two types of behavioural tasks, a multi-arrangement task and a rating task. The stimuli employed in all experiments and tasks were 3D reconstructions of real cells from the olfactory system of *Xenopus laevis* tadpoles. These reconstructions were generated by the team of expert biologists lead by Professor Ivan Manzini at the Department of Animal Physiology and Molecular Biomedicine at Justus Liebig University Giessen. All animal procedures were performed following the guidelines of Laboratory animal research of the Institutional Care and Use Committee of the Justus Liebig University of Giessen (649_M; GI 15/7 Nr. G 2/2019).

The cell reconstructions could belong to different cell types found in the olfactory epithelium of *Xenopus* tadpoles. We selected 30 cell stimuli for our experiments, such that they would be equally distributed into four known cell classes plus one indeterminate set. The four known cell classes were basal cells, dividing cells, olfactory sensory neurons, and supporting cells. Cells were classified as belonging to these classes through imaging investigations performed by the expert biologists. Cells belonging to the indeterminate set were those that could not be reliably classified by the experts into any of the known groups. These indeterminate cells might have been, for example, immature versions of olfactory sensory neurons or supporting cells.

Cell reconstructions were obtained through 3D-microscopy. Specifically, 3D-microscopy images were processed using u-shape3D^22^, a software package to reconstruct and analyse 3D cell morphologies. The reconstructions were created as 3D triangulated mesh models. The mesh models were further processed for rendering and 3D printing using Blender 2.91 software (The Blender Foundation, Amsterdam, NL). We removed non-manifold edges, edges with no length, faces or face corners with no area, and small disconnected elements. Larger disconnected elements were instead manually reconnected to the cell’s main body.

For the multi-arrangement task, the stimuli were 3D printed versions of the cell reconstructions. The reconstructions were 3D printed out of a light, yellow plastic at a scaling factor of 1:1000 (1 μm = 1 cm). For the rating task, we created video renderings of each mesh model. Each video sequence displayed one mesh model rotating back and forth along frontal and vertical axes at approximately 120 degrees/s. Video sequences were all 30 seconds long and were created with MATLAB R2020b software. Mesh models in print-ready “.stl” format and video renderings for all stimuli employed in our experiments will be made available from our open-access data repository (doi: 10.5281/zenodo.14046923).

### Apparatus

Experiments were programmed in Python 3.7.0 and MATLAB R2020b. In the multi-arrangement task, participants were asked to arrange the full set of 3D printed stimulus objects on a workbench (60 x 60 cm). The position of the stimuli arranged on the workbench was captured with high-precision 3D tracking rig^53^ employing hardware and software produced by motion capture company Qualisys (Qualisys AB, Sweden). The stimuli were imaged from multiple angles by 8 tracking cameras (Qualisys Miqus) and 6 video cameras (Qualisys Miqus Video) arranged on a square frame surrounding the workspace (**Figure 1D**). To capture the position of the stimuli, we glued one passive reflective marker on top of each 3D printed cell (**Figure 1C**).

In the rating task, participants viewed stimulus videos on an Asus VG248QE monitor (24’’, resolution = 1920 x 1080 pixel) at 60 Hz, positioned at a distance of 40 cm from the observers (**Figure 1E**). The monitor subtended 67 x 41 degrees of visual angle, and the object size ranged from 1 to 10 degrees of visual angle.

### Procedure

In Experiment 1, naïve participants performed a 1^st^ session of a multi-arrangement task, followed by a rating task, followed by a 2^nd^ session of the multi-arrangement task. In Experiment 2, naïve participants performed only a rating task, which was identical to the rating task performed by participants in Experiment 1. In Experiments 1 and 2, participants were kept naïve to the objects’ origin and were never told that the objects belonged to distinct categories. In Experiment 3, a set of expert participants preformed the same set of tasks and sessions as naïve participants in Experiment 1.

#### Multi-arrangement task

Each session, the experimenter initially positioned the 3D printed stimuli in random order in a circle on the workbench. Stimulus order was pseudo-randomly generated by the experimental script. Participants were asked to arrange the stimuli according to similarity. This meant that similar objects should be placed spatially close together whilst dissimilar objects should be placed far apart. Participants had unlimited time to perform these arrangements. Once a participant was satisfied with the stimulus arrangement, the experimenter recorded the final stimulus positioning measured by the Qualisys tracking system using an ad-hoc script written in Python. The identity of the stimuli was manually annotated by the experimenter off-line.

#### Rating task

Participants performed a computer-based rating task in which they were shown video sequences of rotating 3D stimuli. Each trial, a participant had to rate one of the stimuli along one of eight feature dimensions. Participants rated each stimulus only once for each dimension, for a total of (30×8) 240 trials. Participants had unlimited time to perform these ratings. The order of the stimuli was randomized across participants, but the rating dimensions were queried in the same succession for each stimulus object. Participants made their assessment either by using the mouse to adjust a slider or by typing in a number on the keyboard. To ensure participants understood the rating dimensions, prior to the start of the experiment they were shown exemplary cartoon drawings for each dimension (**Figure 1F**).

### Analyses

We used *MATLAB R2020b* software to analyse the data.

#### Analysis of the multi-arrangement data

First, we used Procrustes analysis to align and visualize the arrangements of the 30 cell models for each participant in each condition. Specifically, we translated and rotated the arrangement data (without rescaling but allowing reflection) so they were best-aligned across participants and sessions. Then, representational dissimilarity matrices for each participant and each session were constructed from the arrangement data by computing the pairwise Euclidean distances between the 30 objects. All analyses were performed using only the upper triangular portions of the dissimilarity matrices, excluding the diagonal.

##### Multi-arrangement task, within-participant agreement

For each participant we computed two dissimilarity matrices, one for the 1^st^ session *RDM_p,s1_* and one for the 2^nd^ session *RDM_p,s2_*. We computed the participant’s average dissimilarity matrix across 1^st^ and 2^nd^ session as:

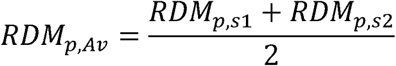

Then we calculated the Pearson correlation between this average dissimilarity matrix and the participant’s 1^st^ and 2^nd^ session dissimilarity matrices:

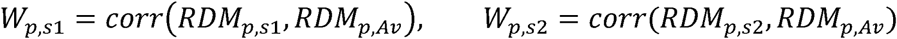

The average of these two correlations was taken as the within-participant agreement:

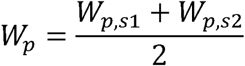

##### Multi-arrangement task, between-participant agreement

We first computed the average dissimilarity matrix across all *N* participants as:

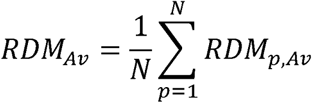

Then, between-participant agreement we calculated as the Pearson correlation between participants’ average dissimilarity matrices *RDM_p,Av_* and the average dissimilarity matrix of all participants:

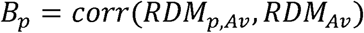

##### Multi-arrangement task, within-session, between-participant agreement

To compute single-session between-participant agreement, we first calculated the single-session average dissimilarity matrices across all *N* participants as:

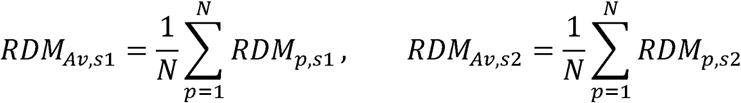

Then, single-session between-participant agreements were computed as:

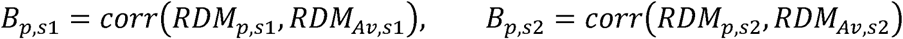

##### Multi-arrangement task, agreement with cell class

The dissimilarity matrix for cell class *RDM_CC_* was constructed as a binary matrix, where cells belonging to the same class had 0 dissimilarity, and cells belonging to different classes had a dissimilarity of 1. The agreement between participant arrangements and cell class was taken as the correlation between the cell class dissimilarity matrix and the average dissimilarity matrix across participants:

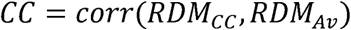

#### Analysis of the rating task data

##### Rating task, between-participant agreement in Experiment 1

For each of the eight feature dimensions *d*, we first computed the average ratings across participants for all 30 stimuli. Between-participant agreement was taken as the correlation between individual participant ratings *Rt_p,d_* and the average rating across participants *Rt_Av,d_*.

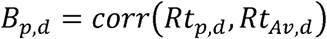

##### Between-task agreement within Experiment 1

We computed dissimilarity matrices for each rating dimension and for every participant *RDM^Rt^_p,d_*. To compute between-task agreement, we then correlated these dissimilarity matrices with the average dissimilarity matrix from the multi-arrangement task:

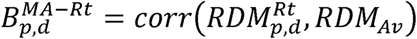

We next fit a linear model regressing individual participant rating dissimilarity matrices onto the average multi-arrangement dissimilarity matrix. The model was defined as:

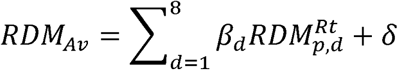

Where *β_d_* are linear coefficients and δ is the intercept.

##### Rating task agreement across Experiments 1 and 2

Between-participant agreement across Experiments 1 and 2 was taken as the correlation between individual participant ratings in Experiment 2, *Rt^E2^_p,d_* and the average rating across participants in Experiment 1:

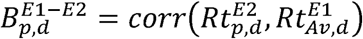

##### Between-task agreement across Experiments 1 and 2

We computed dissimilarity matrices for each rating dimension and for every participant from Experiment 2 *RDM^Rt,E2^_p,d_*. To compute between-task agreement across Experiments 1 and 2, we correlated these dissimilarity matrices with the average dissimilarity matrix from the multi-arrangement task in Experiment 1:

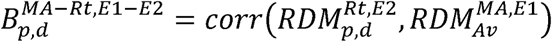

Finally, we used the coefficients fitted above to predict, from the rating data in Experiment 2, the multi-arrangement dissimilarity matrix from Experiment 1.

##### Principal Component Analysis of the rating task data

We assessed the multicollinearity of the rating dimensions by computing the pairwise correlations between all the 8 rating features. We then performed Principle Component Analysis to recover 8 new dimensions, or principal components, each one orthogonal (and thus uncorrelated) to the others, composed of a linear combination of the original features. To prevent biased results, we standardized the scaling of the two dimensions which were assessed by assigning a discrete number (*main body parts* and *number of limbs*) to the same [0 1] range in which the other 6 dimensions were expressed.

##### Agreement between rating data and cell class

To investigate whether rating data aligned with the underlying biological cell class, we computed per-participant RMDs for each PCA dimension, and then fit a linear model regressing individual participant dissimilarity matrices onto the cell class dissimilarity matrix.

#### Classification Analyses

Our final analyses tested whether it was possible to recover the underlying biological cell classes of our stimuli from human perceptual arrangements and judgments of 3D shape. We trained support vector machine classifiers to classify the participant multi-arrangement and rating data into the correct underlying cell class. We performed these analyses on the data from Experiment 1 (naïves) and Experiment 3 (experts). Classification accuracy was computed using the cross-validation procedure described in the Results.

## Author Contributions

**Kira Dehn**: Conceptualization; Methodology; Software; Validation; Formal analysis; Investigation; Data Curation; Writing - Original Draft; Writing - Review & Editing; Visualization, Project administration. **Guido Maiello**: Conceptualization; Methodology; Software; Validation; Formal analysis; Investigation; Resources; Data Curation; Writing - Original Draft; Writing - Review & Editing; Visualization, Supervision, Project administration. **Frieder Hartmann**: Conceptualization; Methodology; Software; Writing - Review & Editing. **Yaniv Morgenstern**: Conceptualization; Methodology; Writing - Review & Editing. **Sara Joy Hawkins**: Conceptualization; Methodology; Resources; Writing - Review & Editing. **Thomas Offner**: Conceptualization; Methodology; Resources; Writing - Review & Editing. **Joshua Walter**: Conceptualization; Methodology; Resources; Writing - Review & Editing. **Thomas Hassenklöver:** Conceptualization; Methodology; Resources; Writing - Review & Editing. **Ivan Manzini:** Conceptualization; Methodology; Resources; Writing - Review & Editing; Funding acquisition. **Roland W. Fleming**: Conceptualization; Methodology; Resources; Writing - Review & Editing; Supervision, Project administration, Funding acquisition

## Data availability

Data and analysis scripts are publicly available from the Zenodo database (doi: 10.5281/zenodo.14046923)

## Acknowledgments

This research was supported by:

The DFG (SFB-TRR-135-TP-C1: “Cardinal Mechanisms of Perception”, and project PA 3723/1-1).

Research Cluster “The Adaptive Mind” funded by the Excellence Program of the Hessian Ministry of Higher Education, Science, Research and Art.

An ERC Advanced Grant (ERC-2022-AdG-101098225 “STUFF”) to RWF.

